# *De novo* Variants in Neurodevelopmental Disorders with Epilepsy

**DOI:** 10.1101/123323

**Authors:** Henrike O. Heyne, Tarjinder Singh, Hannah Stamberger, Rami Abou Jamra, Hande Caglayan, Dana Craiu, Peter De Jonghe, Renzo Guerrini, Katherine L. Helbig, Bobby P. C. Koeleman, Jack A. Kosmicki, Tarja Linnankivi, Patrick May, Hiltrud Muhle, Rikke S. Møller, Bernd A. Neubauer, Aarno Palotie, Manuela Pendziwiat, Pasquale Striano, Sha Tang, Sitao Wu, EuroEPINOMICS RES Consortium, Annapurna Poduri, Yvonne G. Weber, Sarah Weckhuysen, Sanjay M. Sisodiya, Mark Daly, Ingo Helbig, Dennis Lal, Johannes R. Lemke

## Abstract

Epilepsy is a frequent feature of neurodevelopmental disorders (NDD) but little is known about genetic differences between NDD with and without epilepsy. We analyzed *de novo* variants (DNV) in 6753 parent-offspring trios ascertained for different NDD. In the subset of 1942 individuals with NDD with epilepsy, we identified 33 genes with a significant excess of DNV, of which *SNAP25* and *GABRB2* had previously only limited evidence for disease association. Joint analysis of all individuals with NDD also implicated *CACNA1E* as a novel disease gene. Comparing NDD with and without epilepsy, we found missense DNV, DNV in specific genes, age of recruitment and severity of intellectual disability to be associated with epilepsy. We further demonstrate to what extent our results impact current genetic testing as well as treatment, emphasizing the benefit of accurate genetic diagnosis in NDD with epilepsy.

## Introduction

Epilepsies, defined as recurrent, unprovoked seizures, affect about 50 million people worldwide (www.who.int, 03/2017). A significant subset of severe and intractable epilepsies starts in infancy and childhood and poses a major clinical burden to patients, families, and society^1^. Early onset epilepsies are often comorbid with neurodevelopmental disorders (NDD), such as developmental delay, intellectual disability and autism spectrum disorders (DD, ID, ASD)^2–4^, while up to 26% of individuals with NDD have epilepsy, depending on the severity of intellectual impairment^4–6^. Several genes have been implicated in both NDD and epilepsy disorders^7,8^. The epileptic encephalopathies (EE) comprise a heterogeneous group of epilepsy syndromes characterized by frequent and intractable seizures that are thought to contribute to developmental regression^39^. Phenotypic categorisation of clinically-recognizable EE syndromes enabled identification of several associated genes^1,2,10^. However, the phenotypic spectrum of these disease genes was broader than expected^11,12^, ranging from EE (e.g. *SCN1A*^13^, *KCNQ2*^14^) to unspecific NDD with or without epilepsy (e.g. *SCN2A*^15^, *STXBP1*^16^). While clinically distinguishable entities exist, many patients with NDD and epilepsy are not easily classified into EE syndromes^1,12^. Consequently, EE is often used synonymously with NDD with epilepsy^17^. Targeted sequencing of disease-specific gene panels is commonly used in diagnostics of epilepsies^12,18,19^. However, epilepsy gene panel designs of diagnostic laboratories differ substantially in gene content^19^.

Application of a mutational model^18^ to detect enrichment for *de novo* variants (DNV) has proven to be a powerful approach for identification of disease-associated genes in neurodevelopmental disorders including ID, congenital heart disease, schizophrenia and ASD^20–23^. For EE, the currently largest exome-wide DNV burden study comprised 356 parent-offspring trios of two classic EE syndromes (infantile/epileptic spasms, IS and Lennox-Gastaut syndrome, LGS) and revealed seven genes at exome-wide significance^24^. To identify genes that are significantly associated with NDD with epilepsy, we analysed 6753 parent-offspring trios of NDD, focusing on 1942 cases with epilepsy including 529 individuals with epileptic encephalopathy. We compared rates of DNV between EE, NDD with unspecified epilepsies and NDD without epilepsy to identify genetic differences between these phenotypic groups. We further investigated the potential impact of our findings on the design of genetic testing approaches and assessed the extent of therapeutically relevant diagnoses.

## Results

### Description of dataset

We analysed DNV in parent-offspring trios of eight published^7,20,23–27^, one partly published^28^ and three unpublished cohorts of in total 6753 individuals with NDD stratifying for the 1942 cases with epilepsy (Supplementary Table 1, Figure 1, Online Methods). These 1942 patients were ascertained for either EE or NDD with unspecified epilepsy (DD^21^, ASD^11^ with ID and ID^20^). We define those two phenotype groups as NDD_EE_ (n = 529) and NDD_uE_ (n = 1413), respectively. We later compared DNV in NDD with epilepsy (NDD_EE+uE_) to DNV in NDD without epilepsy (NDD_woE_, n = 4811). For genotype-phenotype comparisons, we restricted our analysis to regions that were adequately captured across different capture solutions (see Online Methods). For ASD data from the Simon Simplex Consortium^29^, we included only individuals with IQ < 70 (defined as ID) as different studies have found DNV only associated with low-IQ ASD^6,30^. Individuals with NDD_EE_ were diagnosed with following specific syndromes: IS (n = 243), LGS (n = 145), electrical status epilepticus in sleep (ESES, n = 42), myoclonic-atonic epilepsy (MAE, n = 39), Dravet syndrome (DS, n = 16), unspecified EE (n = 44). Six of eight NDD cohorts (n = 6037) included individuals with as well as without epilepsy^20,23,25–27,31^. Of these, 20.3% of patients had epilepsy. In cohorts with more severe ID, a higher rate of patients had epilepsy (Spearman-Rank correlation, p-value = 0.012, rho = 0.89, Supplementary Figure S2), in line with previous literature^46^. We considered DNV of 1911 healthy siblings of patients with ASD as a control group.

**Figure 1.**
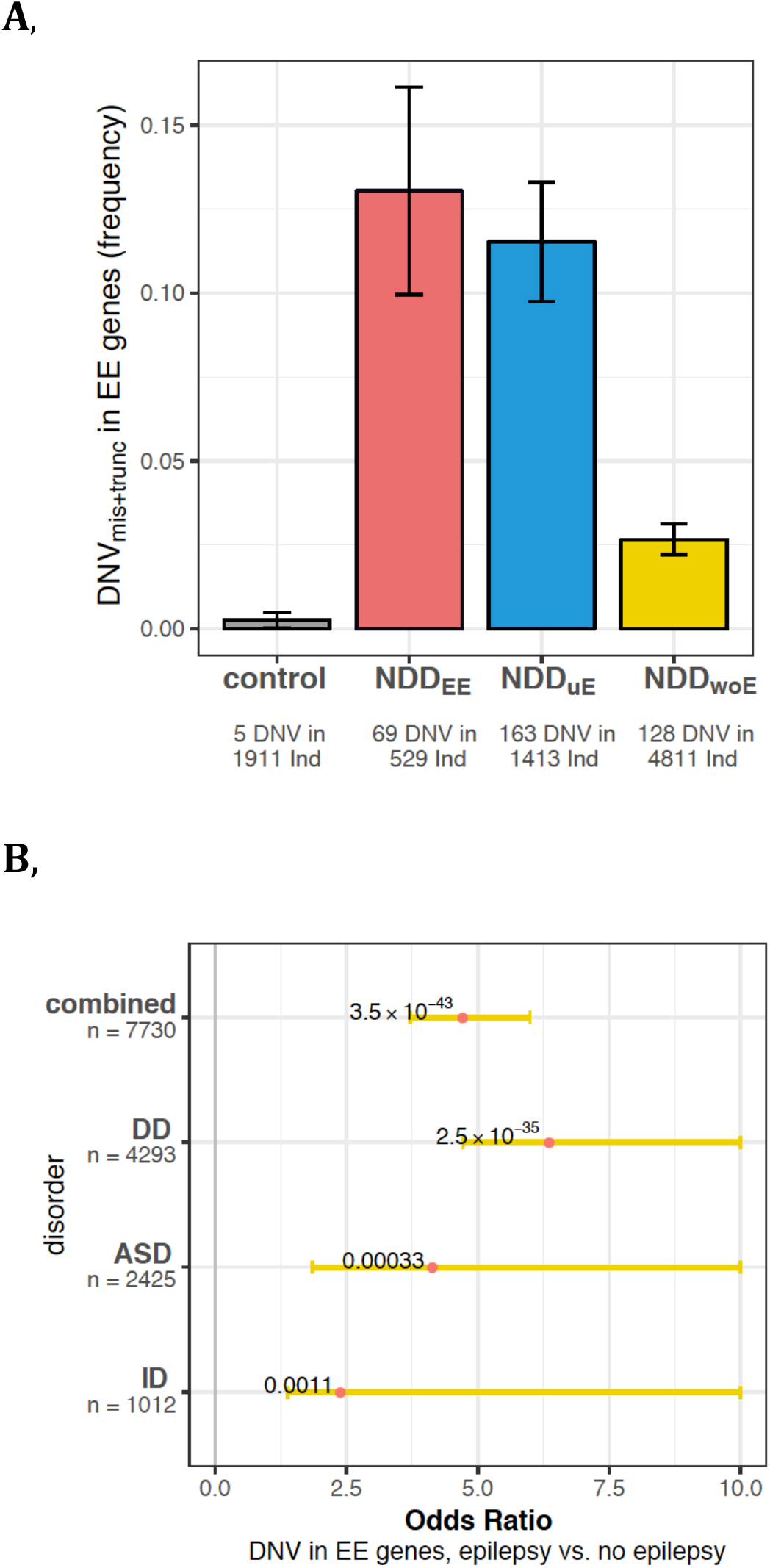
DNV_mis+trunc_ in EE genes in different cohorts of NDD. **A**, The proportion of DNV_mis+trunc_ in EE genes is not significantly different between patients with NDD_EE_ (red) and NDD_uE_ (blue), but higher than NDD_woE_ (yellow) or healthy controls (grey). Cohort size is given as number of individuals (Ind). **B**, Proportion of DNV in EE genes in patients with versus without epilepsy across different NDD (DD, ASD, ID). P-values are plotted next to respective odds ratios (red dots), while 95%-CI are shown as yellow bars (Fisher’s exact test for individual cohorts, Cochran-Mantel-Haenszel test for combined cohorts).

### DNV in known EE genes in patients with different NDD diagnoses

We first compared DNV in known EE genes between NDD_EE_, NDD_uE_, NDD_woE_ and control cohorts. We investigated missense and truncating DNV (DNV_mis+trunc_) in 50 known autosomal dominant or X-linked EE genes (updated list from^19^, Supplementary Table 3). We excluded DNV present in ExAC^32^ to improve power, as these have been shown to confer no risk to childhood-onset NDD on a group level^33^. The frequency of DNV_mis+trunc_ in EE genes was not significantly different between NDD_EE_ (13.0%±3.1, mean, 95%-CI) and NDD_uE_ (11.5%±1.8, mean, 95%-CI, p-value = 0.4, Fisher’s Exact Test, Figure 1A, see Supplementary Figure S2 for individual cohorts), but was significantly greater than in NDD_woE_ (2.7%±0.5, mean, 95%-CI, p-value = 4.4×10^−46^) and in healthy controls (0.3%±0.2, mean, 95%-CI)^20^. Within three different NDD diagnoses (ID, ASD [with and without ID], DD), we detected more DNV in EE genes in individuals with epilepsy than without epilepsy (Cochran-Mantel-Haenszel test, p-value 3.5×10^−43^, common OR 4.6, 95%-CI: 3.7 to 5.9, Figure 2B). This suggests a markedly overlapping genetic spectrum of NDD_EE_ and NDD_uE_. We subsequently performed DNV enrichment analyses on the combined cohort of NDD_EE+uE_.

**Figure 2.**
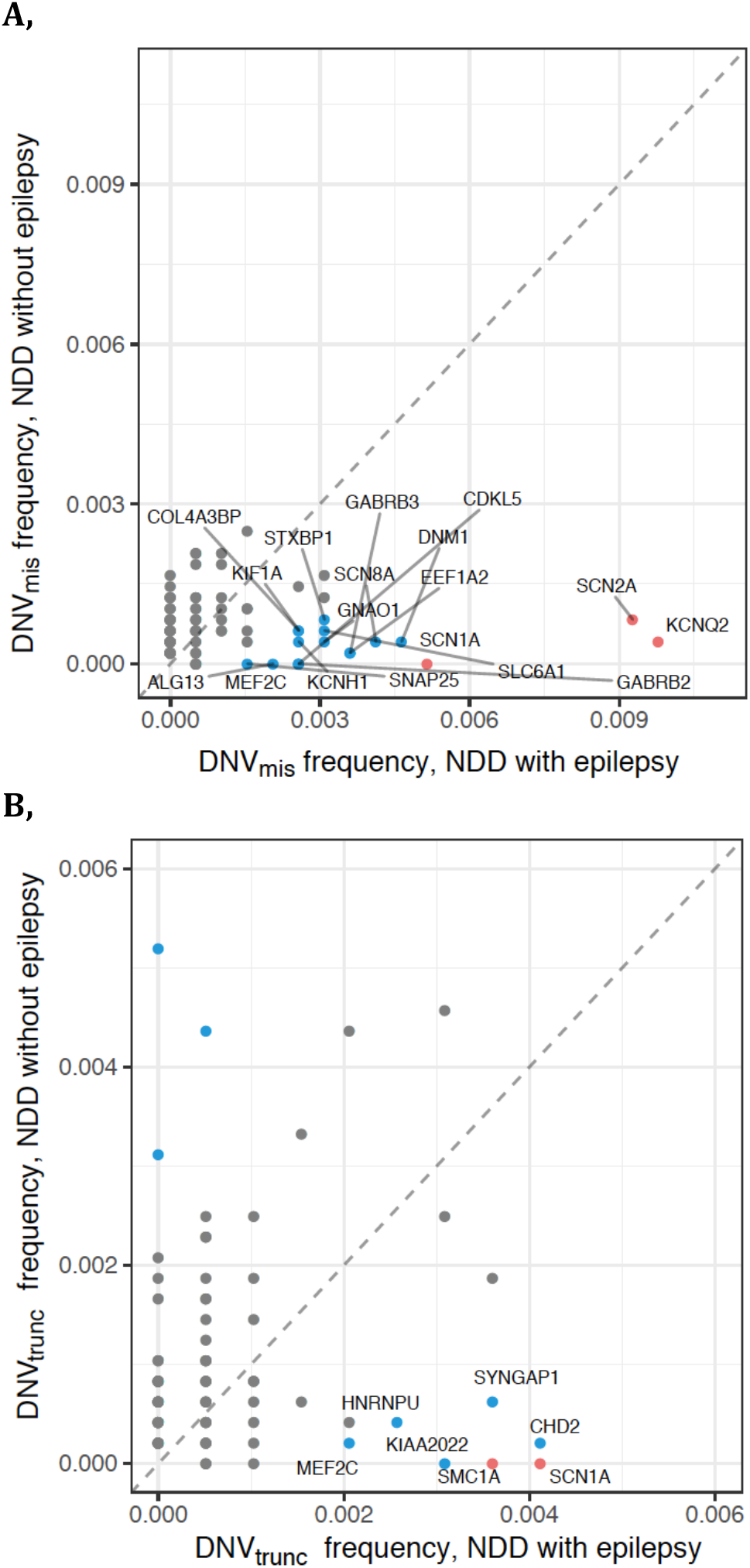
DNV in patients with epilepsy (NDD_EE+uE_) versus without epilepsy (NDD_woE_) in 107 genes with significant DNV burden. **A**, DNV_mis_, **B**, DNV_trunc_. Genes with different frequencies are labeled (method: Fisher’s Exact test, blue: nominal significance, p-value < 0.05, red: significant after correcting for 266 tests). The dotted line represents equal frequency of DNV in NDD with and without epilepsy.

### Discovery of genes with exome-wide DNV burden in NDD with epilepsy

We compared the numbers of DNV in the combined cohort of NDD with epilepsy (NDD_EE+uE_), to the number of DNV expected by a mutational model^30^ revealing global enrichment of truncating (2.3-fold, ptrunc= 1 × 10^−47^, Poisson Exact test, see Online Methods) and missense (1.6-fold, pmis=2 × 10^−33^) but not synonymous DNV (0.6 fold, psyn=1.0). We identified 33 genes with an exome-wide significant burden of DNV_mis+trunc_ (Table 1), of which *KCNQ2* (n=21), *SCN2A* (n=20) and *SCN1A* (n=19) were most frequently mutated. *GABRB2* and *SNAP25* had previously no statistical evidence for disease association (see Supplementary Note). Beyond the 33 genes with exome-wide significant DNV burden, 114 genes had at least two DNV_mis+trunc_ in our cohort (Supplementary Table 6). After DNV enrichment analysis, we again excluded DNV in ExAC^32^ to improve specificity^33^.

**Table 1.** Genes with exome-wide DNV burden in NDD with epilepsy

| Gene | DNV<sub>trunc</sub> |  | DNV<sub>mis</sub> |  | DNV<sub>mis+trunc</sub> |
| --- | --- | --- | --- | --- | --- |
|  | yes | no | yes | no | yes+no |
| KCNQ2 | 0 | 1 | 21 | 3 | 25 |
| SCN2A | 2 | 12 | 18 | 5 | 37 |
| SCN1A | 8 | 0 | 11 | 0 | 19 |
| CHD2 | 9 | 1 | 3 | 2 | 15 |
| SYNGAP1 | 10 | 7 | 1 | 2 | 20 |
| STXBP1 | 4 | 3 | 7 | 5 | 19 |
| SCN8A | 0 | 1 | 10 | 3 | 14 |
| MEF2C | 4 | 1 | 5 | 0 | 10 |
| SLC6A1 | 2 | 1 | 7 | 3 | 13 |
| DNM1 | 0 | 0 | 9 | 2 | 11 |
| EEF1A2 | 0 | 0 | 8 | 3 | 11 |
| CDKL5 | 2 | 0 | 6 | 0 | 8 |
| DYRK1A | 7 | 9 | 0 | 5 | 21 |
| SMC1A | 7 | 0 | 0 | 2 | 9 |
| GABRB3 | 0 | 0 | 7 | 1 | 8 |
| KIAA2022 | 6 | 0 | 0 | 0 | 6 |
| ASXL3 | 6 | 12 | 0 | 0 | 18 |
| WDR45 | 5 | 5 | 1 | 0 | 11 |
| ARID1B | 6 | 28 | 0 | 2 | 36 |
| GNAO1 | 0 | 1 | 6 | 2 | 9 |
| ALG13 | 0 | 0 | 6 | 0 | 6 |
| KCNH1 | 0 | 0 | 6 | 2 | 8 |
| GRIN2B | 0 | 3 | 6 | 9 | 18 |
| HNRNPU | 5 | 2 | 0 | 1 | 8 |
| PURA | 3 | 4 | 2 | 4 | 13 |
| GABRB2 | 0 | 0 | 5 | 1 | 6 |
| COL4A3BP | 0 | 0 | 5 | 4 | 9 |
| MECP2 | 2 | 5 | 3 | 5 | 15 |
| FOXG1 | 2 | 3 | 3 | 3 | 11 |
| ANKRD11 | 4 | 28 | 0 | 2 | 34 |
| SNAP25 | 1 | 0 | 3 | 0 | 4 |
| DDX3X | 3 | 19 | 1 | 11 | 34 |
| IQSEC2 | 3 | 2 | 1 | 3 | 9 |

*\*Genes in order of decreasing numbers of DNV<sub>mis+trunc</sub> in NDD with epilepsy*

Collectively analysing all patients with NDD with or without epilepsy (n = 6753), we found 101 genes with exome-wide DNV burden (Supplementary Table 7). Among these 101 genes five were mutated in at least one individual with EE and at least two other individuals with epilepsy with DNV in the same variant class. Of these, *SMARCA2 DYNC1H1* and *SLC35A2* were formerly associated with NDD with epilepsy. *KCNQ3* had previously limited association with NDD with epilepsy and *CACNA1E* had previously no statistical evidence for disease association (Genes further described in Supplementary Notes).

### Phenotypic, biological and therapeutic properties of genes with DNV burden in NDD with epilepsy

We aimed to explore whether the 33 genes with DNV burden in NDD with epilepsy (NDD_EE+uE_) were associated with specific phenotypes. Analyses of human phenotype ontology^34^ (HPO) terms revealed most significant enrichment in genes associated with “epileptic encephalopathy” (see Online Methods, Supplementary Table 8). After excluding the 529 patients diagnosed with EE from the DNV enrichment analysis, the most significantly enriched HPO term was still “epileptic encephalopathy” (Bonferroni p-value 3.6 × 10^−14^), confirming our previous findings (Figure 1). Per DNV-enriched gene, we plotted distribution of EE phenotypes, sex and seizure phenotypes of generalized, focal, febrile or spasms (Supplementary Figure S6-8).

Since the disease onset of NDD with epilepsy is typically in infancy and early childhood, we evaluated expression levels of the 33 genes with DNV burden in the developing infant brain (expression data: brainspan.org, see Online Methods). At a group level, these genes showed high levels of brain expression (Supplementary Figure S9A). The DNV-enriched genes were also substantially depleted for truncating and missense variants in the ExAC control data (Supplementary Figure S9B, S9C). Genes with at least two DNV in NDD_EE+uE_, but no significant DNV burden showed similar patterns.

We finally evaluated if genes with DNV_mis+trunc_ in NDD with epilepsy were associated with therapy. For each gene, we used criteria from the Centre for Evidence-Based Medicine (CEBM)^35^ to evaluate the evidence for targeted treatments. Five of the 33 DNV-enriched genes (*SCN1A, SCN2A, SCN8A, KCNQ2, MECP2*) had evidence for therapeutic relevance (CEBM Grade of Recommendation A and B, see Online Methods, Supplementary Table 9). These five genes accounted for 28% of all DNV_mis+trunc_ in the significantly implicated genes. Three additional genes (*PTEN, CACNA1A, SLC2A1*) with at least two DNV_mis+trunc_, which were also known disease genes, also had therapeutic relevance according to CEBM criteria. In total 5% (84/1587) of DNV_mis+trunc_ in NDD with epilepsy were in genes with therapeutic consequences. According to the guidelines of the American College of Medical Genetics (ACMG,)^36^ all DNV that are not in ExAC and that are in known disease genes or genes with DNV burden in our dataset are categorized as “likely pathogenic”, while we did not apply all ACMG criteria to individual DNV (see online methods).

### Comparing DNV between NDD with and without epilepsy

We compared frequencies of DNV_mis+trunc_ in NDD with epilepsy (NDD_EE+uE_) to NDD_woE_ across all 107 DNV-enriched genes (logistic regression, see Online Methods). Increasing age at time of recruitment increased likelihood of epilepsy (three-year OR 1.11, 95%-CI 1.04 to 1.18, p-value = 3×10^−3^, individual genes in Supplementary Figure S5). Sex was not associated with epilepsy status (p-value = 0.5). Individuals with DNV_mis_ were more likely to have epilepsy than individuals with DNV_trunc_ (Figure 2, ORmis 2.1, 95%-CI 1.6 to 2.8, p-value 2×10^−7^). In line with previous reports^15^, we observed this pattern on a single gene level for *SCN2A* (Firth regression, ORmis 23.5, 95%-CI 3.8 to 277, p-value 0.0003, Table 1). Confirming previous findings^24,37^, DNV in ion channel genes were associated with epilepsy (OR 6.0, 95%-CI 3.9 to 9.2, p-value 1×10^−16^). 83% (110/133) of DNV in ion channel genes were DNV_mis_. However, in the subset of 910 DNV not in ion channel genes, DNV_mis_ were still associated with epilepsy (OR 1.5, p-value 0.005, 95%-CI 1.1 to 2.1), implying that the effect of DNV_mis_ on epilepsy was not entirely driven by ion channel genes. We observed a higher rate of DNV_mis_ in NDD_EE_ than in NDD_uE_, though only with nominal significance (Fisher’s exact test, OR 1.8, 95%-CI 1.04 to 3.4, p-value 0.03, Supplementary Figure S10B). Four genes were more frequently mutated in NDD with epilepsy (NDD_EE+uE_) than NDD_woE_ (Fisher’s Exact Test, Figure 2A/2B, Table 1, Supplementary Table 10). With the exception of *SCN1A*, frequencies of DNV were not significantly different per gene between NDD_EE_ and NDD_uE_ for DNV_mis_ or DNV_trunc_ (Supplementary Figure S10, Supplementary Table 11).

### Evaluation of diagnostic gene panels for epilepsy disorders

Targeted sequencing of disease-specific gene panels is widely employed in diagnostics of epilepsies^18,19^. We compared our results to 24 diagnostic panels for epilepsy or EE (see Online Methods, full list in Supplementary Table 12). In total, the 24 different panels covered 358 unique genes (81.5 ± 8.8 genes per panel, mean ± sd). Applying these 24 diagnostic panels on our data set would only have detected on average 59% of DNV_mis+trunc_ in the 33 DNV-enriched genes (Supplementary Figure S11). However, similar to most other research studies involving clinical WES^7^, we cannot fully assess the extent of potential pre-screening. We investigated whether genes in the 24 panels had some evidence for disease association given the following features that we (and others^23,33^) observed in genes with DNV burden in NDD: depletion for truncating and missense variants in ExAC^32^ controls as well as brain expression (Online Methods, Supplementary Figure S9). We restricted this analysis to autosomal dominant and X-linked acting panel genes (ndominant+x-linked = 191, Supplementary Table 13). 95% (52/55) of panel genes that had two or more DNV_mis+trunc_ in our study were both constraint and brain-expressed. However, only 63% (86/136) of panel genes with one or less DNV_mis+trunc_ in our study were constraint and brain-expressed (Fisher’s exact test, OR 10.2, 95%-CI 3.0 to 53.0, p-value 2.3 × 10^−6^). We applied evidence of disease association as defined by the ClinGen Gene Curation Workgroup^38^, to those 50 panel genes lacking two of the criteria DNV/brain expression/constraint. We found that ten of the 50 genes had no, eight had limited and seven had conflicting published evidence for disease association (Supplementary Table 14). Thirteen genes showed moderate, strong or definitive evidence for association to entities where neither NDD nor epilepsy were major features which may partly be explained by a panel design containing genes associated with diseases beyond the spectrum of NDD (for further details see Online Methods and Supplementary Figure S11).

## Discussion

In this study, we systematically investigated DNV in NDD with and without epilepsy. In NDD with epilepsy, we could hardly distinguish individuals ascertained for epileptic encephalopathy and NDD with unspecified epilepsy on a genetic level. Thus, we conclude that these phenotype groups share a spectrum of disease genes predominantly including genes initially reported as EE genes. We identified 33 genes with DNV burden in NDD with epilepsy, of which the majority was expressed in the infant brain and depleted for functional variation in ExAC^32^, as previously described for NDD genes^23,33^. We report statistically robust disease association for *SNAP25, GABRB2* and *CACNA1E*, which was previously lacking (Supplementary Notes).

We found, that individuals with DNV_mis_ were generally more likely to have epilepsy than individuals with DNV_trunc_. This association was largely driven by ion channel genes, which confirms longstanding statements that many epilepsy disorders act as channelopathies^2,37,24^. Heterozygous DNV_mis_ have been shown to cause epilepsy via dominant negative (e.g. *KCNQ2*^39^) or gain-of-function (e.g. *SCN8A*^40^) effects on ion channels. On the individual gene level, missense variants in *SCN2A*^15^ and *SCN8A*^41^ were more strongly implicated in epilepsy than protein truncating variants, which we statistically confirm for *SCN2A*. Yet, we found that DNV_mis_ were also associated with epilepsy independent of ion channel genes. This may imply that alteration of protein function quantitatively plays a larger role than haploinsufficiency^42^ in the pathophysiology of NDD with epilepsy compared to NDD without epilepsy. We found multiple gene sets enriched for DNV_mis_ in epilepsy compared to no epilepsy (see Supplementary Note). The majority was related to ion channels, while others related to neuronal cells (e.g. axon part, synaptic transmission). However, biological interpretation should be done with caution given that previous studies have found that many of these gene sets share a large number of underlying genes^22^ and gene annotations are biased^43^. We further replicate a previous finding that the rate of epilepsy was correlated with severity of intellectual disability^4–6^, implying that brain function could contribute to epileptogenesis or genetic variants cause both epilepsy and NDD. Alternatively, severe epileptic activity may also damage brain function and thereby contribute to NDD, which constitutes the original definition of EE^9,17^. This is supported by many cases of clinical regression after onset of epilepsy and improvement of NDD through seizure control.

In NDD with epilepsy we found no genetic differences between unspecified epilepsy and EE, with the exception of *SCN1A* (Supplementary Note). Phenotypic heterogeneity has been described for the majority of EE genes^1,11^, i.e. variants in the same gene could lead to a spectrum of different phenotypes. Due to pleiotropy, individuals that carry a pathogenic DNV in an EE gene and fulfil diagnostic criteria of EE may also be eligible for another NDD diagnosis and thus by chance be assigned to an ASD, DD or ID and not an EE screening cohort. In line with this hypothesis, we found typically EE-associated seizure types (e.g. epileptic spasms) in cohorts with unspecified epilepsy. Some of the diagnostic criteria for EE^1,10^ may present ambiguously, leading to uncertainty in terminology^17^. Thus, 43% (21/49) of individuals diagnosed with EE in the Epi4K-E2^24^ study initially presented with DD prior to seizure onset conflicting with the original definition of EE^3,17^. Clear phenotypic distinction between encephalopathic versus non-encephalopathic epilepsies may therefore be difficult. Accordingly, mechanisms that result in an encephalopathic course of a genetic NDD remain elusive.

Restricting DNA sequencing or DNA sequence analysis to panels of known disease genes is widely used in diagnosis of genetic diseases including epilepsy (^19^, 100,000 genomes project [www.genomicsengland.co.uk]). We confirmed that epilepsy gene panels from diagnostic laboratories differ substantially in gene content^18^ with at least 25 genes with low evidence for disease association (ClinGen criteria^38^). Statistically not robust gene-disease associations occasionally resulted in false-positive reports of causality posing challenges for correct diagnosis in research and clinical settings^11,44^. Our data provide grounds for replacing genes with limited evidence by genes with higher evidence in the design of gene panels for NDD with epilepsy.

Therapeutic approaches, tailored to the patient’s underlying genetic variant, have successfully been applied for several EE^2^ including treatment with ezogabine in *KCNQ2* encephalopathy^45^ or ketogenic diet in *SLC2A1*-related disorders^46^. 5% of DNV_mis+trunc_ in our study were in eight genes (Supplementary Table 9) for which we could confirm therapeutic consequences with established evidence-based medicine criteria^35^. This finding reinforces the urgency of making a genetic diagnosis in NDD with epilepsy. We expect that with increasing understanding of the underlying pathomechanisms, the group of genetic epilepsies with relevant therapeutic consequences will continue to grow.

## Online Methods

### Patient cohorts

For this study, we ascertained 8,529 patients with the following neurodevelopmental disorders (NDD): developmental delay (DD^31^, n=4293), autism spectrum disorder (ASD^20^, n=2508), epileptic encephalopathy^24,28^ (NDD_EE_, n=529), intellectual disability^23,25-27^ (ID, n=1035), and epilepsy with NDD^28^ (n=164). From this cohort, we selected 6753 individuals, for which the presence or absence of epilepsy was ascertained and of whom ca. 88% had ID (based on assumption of 81.7% ID in the DDD study^47^, 89.8% ID in a diagnostic cohort from AmbryGenetics^28^ and 100% ID in all other cohorts.) Among individuals with ASD who were phenotyped within the Simon Simplex Consortium^29^, we restricted our analysis to patients with ID (IQ < 70) as it has been shown that DNV play only a minor role, in normal IQ ASD^6,30^. Previously sequenced trios (n = 1911), from unaffected siblings of a child with ASD^20,29^, served as control trios. For our main analyses, we stratified this combined cohort of patients with NDD for patients comorbid or primarily diagnosed with epilepsy (NDD_EE+uE_, n=19 42)^20,29^. Two EE cohorts and one ID cohort comprising a combined 144 patients were not previously published; one cohort was only partly published (see Supplementary Table 1). Medical doctors, mostly clinical geneticists, but also neurologists, paediatricians and for ASD^29^ some primary care physicians reported out phenotypes, including presence of epilepsy, in all patients. Our analysis is based on the assumption that medical professionals are sufficiently qualified to diagnose the presence or absence of epilepsy correctly.

### Subphenotypes

We obtained information on specific EE syndromes on 98% of 518/529 individuals with NDD_EE_ (see main text). We obtained specific seizure types (febrile, focal, spasms, generalized) for 55% (140/256) and age of seizure onset for 30% (77/256) of individuals with DNV_mis+trunc_ in genes with DNV burden in NDD_EE+uE_. (See Supplementary Figure S5 and S6). We did not obtain EEG data per patient. Some patients may have developed epilepsy after inclusion in the study, so we ascertained age at recruitment, that we obtained for 94% (1087/1157) of all individuals with NDD with DNV_mis+trunc_ in DNV-enriched genes (median age at recruitment: 74.8 months). We obtained age of seizure onset for 30% (77/256) of individuals with epilepsy and DNV_mis+trunc_ in DNV-enriched genes (Supplementary Figure S5). We identified 30 individuals with potentially epilepsy-relevant brain malformations (abnormalities of neuronal migration, structural abnormalities of corpus callosum, midbrain, brainstem as schiz-, megal-, holoprosencephaly) in individuals with DNV_mis+trunc_ in DNV-enriched genes (29 from DDD^29^, 1 from Hamdan *et al*.^7^). 11 of them (37%) also had seizures.

### Whole exome sequencing of parent-patient trios

In all cohorts, both patients and their unaffected parents underwent whole exome sequencing (WES). Variants that were not present in either parent were considered *de novo* variants (DNV). 1942 individuals with NDD with epilepsy (NDD_EE+uE_) had 1687 DNV_mis_ and 396 DNV_trunc_ (i.e. stopgain, frameshift, essential splice site). 4811 individuals with NDD_woE_ had 4227 DNV_mis_ and 1120 DNV_trunc_ (Supplementary Table 2, for individual cohorts see Supplementary Figure S3). The study was approved by the ethics committee of the University of Leipzig (224/16-ek, 402/16-ek) and additional local ethics committees. A list of all published and unpublished cohorts used in this paper can be found in Supplementary Table 1.

### Sequencing pipelines of previously unpublished/partly published cohorts (cohorts 8-11)

Libraries were prepared from parents’ and patients’ DNA, exome captured and sequenced on Illumina sequencers. Raw data was processed and technically filtered with established pipelines at the respective academic or diagnostic laboratories. DNV data from all cohorts was re-annotated for this study (see below). Specific pipelines of cohorts 10 to 14 are described below.

### Cohort 8 (Ambry Genetics)

Diagnostic WES was performed on parent-offspring trios at Ambry Genetics (Aliso Viejo, CA) in 216 individuals with a history of seizures who have been previously described^28^. Genomic DNA extraction, exome library preparation, sequencing, bioinformatics pipeline, and data analyses were performed as previously described^48^. Briefly, samples were prepared and sequenced using paired-end, 100 cycle chemistry on the Illumina HiSeq 2500 sequencer. Exome enrichment was performed using either the SureSelect Target Enrichment System 3.0 (Agilent Technologies) or SeqCap EZ VCRome 2.0 (Roche NimblGen). The sequencing reads were aligned to human reference genome (GRCh37) and variants were called by using CASAVA software (Illumina). The following variants filters were applied to generate a list of high confident de novo variant calls: 1) mutation base coverage >= 20x in all members of the trio; 2) heterozygous read ratio in probands >30% and <80%; 3) heterozygous read ratio in parents <2%; 4) genotype quality cutoffs SNV > 100 and indels > 300 and 5) exclusion of known sequencing artefacts (based on Ambry Genetics’ internal databases).

### Cohorts 9 (EuroEPINOMICS RES) and 10 (DFG atypical EE)

Exonic and adjacent intronic sequences were enriched from genomic DNA using the NimbleGen SeqCap EZ Human Exome Library v2.0 enrichment kit. WES was performed using a 100bp paired-end read protocol due to the manufacturer’s recommendations on an Illumina HiSeq2000 sequencer by the Cologne Center for Genomics (CCG), Cologne, Germany. Reads were mapped on the human hg19 reference genome (bwa-aln software, bio-bwa.sourceforge.net/). The UnifiedGenotyper (GATK, www.broadinstitute.org/gatk/) and Mpileup (Samtools, http://samtools.sourceforge.net/) software were used to call variants. The paired sample feature from the DeNovoGear software was further used to examine potential de novo mutations in twin pairs. Data analysis and filtering of mapped target sequences was performed with the ‘Varbank’ exome and genome analysis pipeline v.2.1 (unpublished; https://varbank.ccg.uni-koeln.de). In particular, we filtered for high-quality (coverage of more than six reads, fraction of allele carrying reads at least 25%, a minimum genotype quality score of 10, VQSLOD greater than-8) and rare (Caucasian population allele frequency < 0.5%) variations on targeted regions + flanking 100bp. In order to exclude pipeline specific artifacts, we also filtered against an in-house cohort of variations, which were created with the same analysis pipeline.

### Cohort 11 (University of Leipzig)

Exome capture was carried out with Illumina’s Nextera Rapid Capture Exome Kit (Illumina, Inc., San Diego, CA, USA). WES was on an NextSeq500 or HiSeq4000 sequencer (Illumina, Inc.) to 2 × 150bp reads at the Centogene AG, Rostock, Germany. Raw sequencing reads were converted to standard fastq format using bcl2fastq software 2.17.1.14 (Illumina, Inc.), and fed to a pipeline at Centogene AG based on the 1000 Genomes Project (1000G) data analysis pipeline and GATK best practice recommendations. Sequencing reads were aligned to the GRCh37 (hg19) build of the human reference genome using bwa-mem (bio-bwa.sourceforge.net/). In addition to GATK HaplotypeCaller (www.broadinstitute.org/gatk/)), variant calling was performed with freebayes (https://github.com/ekg/freebayes) and samtools (http://samtools.sourceforge.net/). Quality filtering of sequencing reads in both parents and children was done according to the following criteria: read depth > 20, quality > 500, frequency of alternative allele between 30 and 70% for the child and not present in the parents, frequency < 1% in internal database, variant called by at least two different genotype callers.

### False positive rates of DNV

In cohorts 1 to 4, all DNV were validated by Sanger sequencing to eliminate false positive calls. In cohorts 5 to 7, through random selection of variants for Sanger validation, the false positive rate was estimated to be approximately 1.4% and < 5 %, respectively. In the clinical cohorts 8 to 11, variants defined as variants worth reporting back to patients (variants of unknown significance or [likely] pathogenic) are normally validated by Sanger sequencing. With this experience, false discovery rates in these cohorts were estimated to be < 5% (personal communications).

### Annotation and Filtering

DNV files were generated and quality-filtered by the individual groups. All DNV were reannotated with the following pipeline. Variants were annotated with Ensembl’s Variant Effect Predictor (http://grch37.ensembl.org/Homo_sapiens/Tools/VEP) of version 82 using database 83 of GRCh37 as reference genome. Per variant, the transcript with the most severe impact, as predicted by VEP, was selected for further analyses. The decreasing order of variant impacts was HIGH, MODERATE, MODIFIER, LOW. Only protein - altering DNV (DNV_mis_ or DNV_trunc_ [premature stop codon, essential splice site, frameshift]) were included in further analyses. Synonymous DNV (DNV_syn_) were analysed as a negative control, as most DNV_syn_ have no effect on amino acid sequence in the protein. Variants that were present in ExAC^32^, an aggregation of 60,706 exome sequences from adult individuals without severe childhood-onset diseases, were excluded after DNV enrichment, as these have been shown to convey no detectable risk to NDD on a group level^33^. For DNV rates per cohort see Supplementary Figure S2. We did not investigate pathogenicity of individual DNV according to the guidelines of the American College of Medical Genetics (ACMG). However, ACMG criteria PS2 (de novo occurrence, with maternity and paternity confirmed) and PM2 (absence from controls) apply to all DNV in our cohort. The combination of PS2 and PM2 classifies a variant as at least “likely pathogenic”. ACMG criteria are only applicable to variants in disease associated genes^36^. Therefore, all DNV in known disease genes and genes with genome-wide DNV burden in our dataset are presumed likely pathogenic DNV.

### Harmonization of different cohorts

The core analysis of our study is the enrichment of DNV_mis+trunc_ compared to expectation by a mutational model in individuals with NDDEE+nsE. For this analysis, we were conservative in assuming that every gene was well captured across all cohorts. However, when comparing DNV burden across different phenotypes we aimed to separate technical from biological differences with the following methods. In exome sequencing, different capture solutions capture specific exonic regions with different efficiencies. These differences have shown to be quite stable within and across different samples of the same capture kits^49^. We therefore generated a list of exons that displayed consistent high coverage across different capture solutions. We collected published and internal data aiming for the highest possible variety of capture kits using 3,000 samples of 5 different capture kits, including NimbleGen SeqCap v2 and v3, Agilent SureSelect v2, v3, and v5). We generated a list of exons where at least 80% of all samples had at least 10x coverage. We excluded the oldest capture kits before calculating the high coverage exons as well as excluding the two oldest cohorts^26,27^ from our list of DNV. Restricting to high coverage regions resulted in a loss of ca. 11% of DNV in DNV-enriched genes. We consequently performed all genotype phenotype comparisons across cohorts (Figures 1A, 2, Supplementary Figures S6-10) with this restricted DNV set. Further, we compared the frequency of DNV_syn_ across all cohorts and excluded cohorts of which DNV_syn_ were not available. In the subset of DNV in high coverage exons, rates of supposedly neutral DNV_syn_ were not different between individuals with and without epilepsy (Poisson Exact test, p-value = 0.48, RR=0.99), NDD_uE_ and NDD_EE_ (p-value = 0.65, RR= 0.94) or NDD and controls (p-value = 0.58, RR=0.99). The frequency of DNV_mis+trunc_ was also not different between individuals with and without epilepsy (p-value=0.5, RR=1.02). Our chances to identify DNV_mis+trunc_ in EE genes in the epilepsy cohort were therefore not inflated by a higher baseline rate of DNV_mis+trunc_ in comparison to NDD_woE_. We reannotated all DNV in the same way as described above.

### Statistical analysis

All statistical analyses were done with the R programming language (www.r-project.org). Fisher’s Exact Test for Count Data, Wilcoxon rank sum test, Poisson Exact Test, Cochran-Mantel-Haenszel test, logistic regression, Firth regression, Spearman correlation, Welch two-sided t-test and calculation of empirical p-values were performed as referenced in the results. For datasets assumed to be normally distributed after visual inspection, mean and standard deviation (sd) are written as mean ± sd. When performing Poisson Exact Tests, we reported effect size as rate ratio (RR), which is the quotient of the two rates compared in the test. For Fisher’s Exact Test and logistic regression analyses, we reported odds ratios (OR). 95% confidence intervals were abbreviated as 95%-CI. The R code used to perform the statistical analyses and figures is available upon request.

### DNV enrichment analyses

To identify genes with a significant DNV burden, we compared numbers of observed with numbers of expected missense, truncating and synonymous DNV per gene using an established framework of gene-specific mutation rates^30^. The analysis was done with the R package denovolyzer^50^, that compares observed versus expected DNV using a Poisson Exact test. We corrected the obtained p-values with the Bonferroni method for the number of genes for which gene specific mutation rates^30^ were available (n= 18225) and six tests resulting in a p-value significance threshold of 5×10^−7^. Genes that passed that significance threshold for either missense, truncating or both missense plus truncating DNV were considered genes with an exome-wide DNV burden. To compare DNV between disease groups, DNV enrichment analyses were carried out in the cohort of all patients with NDD (n=6753) as well as in patients with epilepsy (NDD_EE+uE_, n=1942) and without epilepsy (NDD_woE_, n=4811), but only genes with a DNV_mis+trunc_ burden in the NDD with epilepsy cohort and the combined NDD cohort were reported.

### HPO enrichment analyses

Significantly enriched Human phenotype ontology (HPO) terms were computed with the R package of g:Profiler^34^, using ordered enrichment analysis on significance-ranked proteins (see Supplementary Table 8). Different gene sets were queried using the background gene set of all 18225 genes for which gene specific mutation rates were available^30^. Only terms that were statistically significant with a Bonferroni corrected p-value < 0.01 were reported, as our negative controls (genes with at least two DNV_mis+trunc_ in healthy control) were not enriched for any functional categories below this p-value.

### Therapeutic relevance

To assess if DNV in our cohort were in genes of therapeutic relevance, we searched the literature for treatment recommendations for all established disease genes with at least two DNV_mis+trunc_ in our NDD with epilepsy cohort. We rated the publications with the standardized score of the Oxford Centre for Evidence-Based Medicine^35^. We only reported and considered genes for which at least one treatment recommendation achieved level of evidence of II or higher. For a list of all genes and levels of evidence see Supplementary Table 9.

### Acquisition and processing of brain gene expression data

We downloaded the Developmental Transcriptome dataset of ‘BrainSpan: Atlas of the Developing Human Brain’ (www.brainspan.org, funded by ARRA Awards 1RC2MH089921-01, 1RC2MH090047-01, and 1RC2MH089929-01, 2011). The atlas includes RNA sequencing data generated from tissue samples of developing postmortem brains of neurologically unremarkable donors covering 8 to 16 brain structures. We extracted brain expression data from the 5 donors that were infants aged 0 to 12 months. Per gene, we obtained the median RPKM value of all infant individuals and across brain regions. In all calculations and figures gene expression values are displayed as median (log2 + 1)-transformed RPKM values. We defined infant brain gene expression as median (log2 + 1)-transformed RPKM value > 1. More details about tissue acquisition and sequencing methodology can be found in the BrainSpan website’s documentation.

### Evaluation of genes’ intolerance to protein altering variants

We assessed individual gene tolerance to truncating or missense variants in the general population with the pLI score (probability of being loss-of-function intolerant) and missense z-score. These scores indicate depletion of truncating and missense variants in ExAC^32^ (60,70 6 individuals without childhood onset diseases), respectively. We used gene constraint cut-offs >0.9 for pLI and >3.09 for missense-z scores as recommended by the score developers^32^. We calculated empirical p-values to evaluate if pLI scores of exome-wide and nominally DNV-enriched genes were significantly higher compared to pLI scores of random gene sets as described in^23^. Briefly, we computed the expected pLI for a given gene set with size n by randomly drawing 1,000,000 gene sets with size n from the total 18,225 pLI annotated genes. We computed, how many times the median pLI score of randomly sampled gene sets would exceed the median pLI of the gene set under investigation. To that number we added 1 and divided by the number of total samplings +1 to obtain the empirical p-value.

### Comparing DNV in NDD_EE_, NDD_uE_ and NDD_woE_

We investigated DNV_mis+trunc_ in NDD_EE+uE+woE_ across all 107 genes that were DNV-enriched in NDD_EE+uE_, NDD_woE_ and/or NDD_EE+uE+woE_. We restricted our analysis to DNV not in ExAC^23^ and in high coverage regions. To investigate, if age at time of recruitment, sex or variant class (DNV_mis_/DNV_trunc_) influenced the presence of epilepsy, we tested them as covariates in a logistic regression model with epilepsy as response variable. We aimed to explore, whether DNV in NDD with epilepsy might be associated with ion channels compared to NDD without epilepsy, as it is a long-established hypothesis, that many epilepsies are channelopathies^37^. We extracted a comprehensive gene set of 237 known ion channel genes from 1766 previously described^22^ curated gene sets derived from public pathway databases and publications (see Supplementary Note). To investigate if ion channel genes were associated with epilepsy we included annotation as ion channel gene as a categorical predictor in the logistic regression model. We used Firth regression to assess the effect of variant class on the presence of epilepsy for individual genes. We used Fisher’s Exact test to compare frequencies of DNV per gene between phenotype groups. To account for multiple testing, we corrected p-values for the number of tests performed (Bonferroni method).

### Diagnostic gene panels for epileptic encephalopathy/ comprehensive epilepsy from 24 academic/ commercial providers

We set out to compare our results to diagnostic gene panels for epileptic encephalopathy of international commercial and academic providers. We searched the Genetic Testing Registry (GTR)^51^ of NCBI (date: 01/2017) for providers of tests for “Epileptic encephalopathy, childhood-onset” and identified 16 diagnostic epilepsy panels. We excluded 3 panels with < 20 or > 200 genes and added 11 additional diagnostic providers not registered at GTR to evaluate 24 diagnostic panels targeting epilepsy in general (n=11) or EE specifically (n=13). The gene content covered in each of the 24 gene panels can be found in Supplementary Table 11. Gene lists were freely available for download at the respective providers’ websites. For each of the 33 genes with DNV burden in NDD with epilepsy, we calculated to what proportion they were included in 24 commercial or academic providers of gene panels for epileptic encephalopathy/comprehensive epilepsy. For each gene, we then multiplied the percentage of inclusion in any of the 24 panels by the total number of DNV_mis+trunc_ of that gene in the cohort of 1942 individuals with NDD_EE+uE_.

We investigated if there were genes in the 24 diagnostic gene panels without evidence for implication in NDD with epilepsy. We focused on 191 dominant or X-linked panel genes (listed in Supplementary Table 14). We tested these genes for three criteria of association with NDD with epilepsy: Firstly, if genes had at least two DNV_mis+trunc_ in our study; secondly, whether genes were expressed in the infant brain defined by a median RPKM of all samples and brain regions > 1; thirdly, whether genes had a pLI > 0.9 or missense z-score > 3.09 indicating intolerance to truncating or missense variants^32^. We intersected these lists to nominate genes that did not display features of DNV-enriched genes in this study. On these genes we applied ClinGen criteria^38^ for gene-disease association.

### Data availability

The authors declare that all data used for computing results supporting the findings of this study are available within the paper and its supplementary information files. Raw sequencing data of published cohorts are referenced at the respective publications. Raw sequencing data of cohort EuroEPINOMICS RES have been deposited in the European Genome-phenome Archive (EGA) with the accession code EGAS00001000048 (https://www.ebi.ac.uk/ega/datasets/EGAD00001000021). Raw sequencing data of cohort 10 (DFG atypical EE) will be deposited in a public repository after finalization of the individual project.

## Acknowledgements

We like to thank all patients and their families who participated in this study, as well as the teams who were involved in recruiting patients, samples and data at the respective study sites. We thank Lisenka Vissers and Christian Gilissen for epilepsy and age phenotypes from the cohort of Lelieveld *et al*, 2016 and Jeremy McRae for useful discussions on the DDD cohort (McRae *et al*, 2016). We thank Johannes Krause for support in figure design and helpful discussions. We are grateful to members of ATGU and the Institute for Human Genetics in Leipzig for helpful contributions. Supported by the Eurocores program EuroEPINOMICS, the Fund for Scientific Research Flanders (FWO), the International Coordination Action (ICA) grant G0E8614N, and the University of Antwerp (research fund). HOH was supported by stipends from the Federal Ministry of Education and Research (BMBF), Germany, FKZ: 01EO1501 and the German Research Foundation (DFG): HE7987/1-1. HS is PhD fellow of the Fund for Scientific Research Flanders (1125416N). IH and YGW were supported by DFG grants WE4896/3-1 and HE5415/6-1. RG received funding through the EU 7^th^ Framework Program (FP7) under the project DESIRE grant N602531.

The DDD study presents independent research commissioned by the Health Innovation Challenge Fund [grant number HICF-1009-003], a parallel funding partnership between the Wellcome Trust and the Department of Health, and the Wellcome Trust Sanger Institute [grant number WT098051]. The views expressed in this publication are those of the author(s) and not necessarily those of the Wellcome Trust or the Department of Health. The study has UK Research Ethics Committee approval (10/H0305/83, granted by the Cambridge South REC, and GEN/284/12 granted by the Republic of Ireland REC). The research team acknowledges the support of the National Institute for Health Research, through the Comprehensive Clinical Research Network.

